# Stiffness of Nanoparticulate Mineralized Collagen Scaffolds Triggers Osteogenesis via Mechanotransduction and Canonical Wnt Signaling

**DOI:** 10.1101/2020.03.09.982231

**Authors:** Qi Zhou, Shengyu Lyu, Anthony A. Bertrand, Allison C. Hu, Candace H. Chan, Xiaoyan Ren, Marley J. Dewey, Aleczandria S. Tiffany, Brendan A.C. Harley, Justine C. Lee

## Abstract

The ability of the extracellular matrix (ECM) to instruct progenitor cell differentiation has generated excitement for the development of materials-based regenerative solutions. We previously described a nanoparticulate mineralized collagen glycosaminoglycan (MC-GAG) material capable of inducing *in vivo* skull regeneration approaching 60% of the biomechanical properties of native calvarium without exogenous growth factors or *ex vivo* progenitor cell-priming, suggesting promise as a first-generation material for skull regeneration. Here, we evaluated the contribution of titrating stiffness to osteogenicity by comparing non-crosslinked (NX-MC) and crosslinked (MC) forms of MC-GAG. While both materials were osteogenic, MC demonstrated an increased expression of osteogenic markers and mineralization compared to NX-MC. Both materials were capable of autogenously activating the canonical bone morphogenetic protein receptor (BMPR) signaling pathway with phosphorylation of Smad1/5 (small mothers against decapentaplegic-1/5). However, unlike NX-MC, hMSCs cultured on MC demonstrated significant elevations in the major mechanotransduction mediators YAP (Yes-associated protein) and TAZ (transcriptional coactivator with PDZ-binding motif) expression coincident with β-catenin activation in the canonical Wnt signaling pathway. Inhibition of YAP/TAZ activation reduced osteogenic marker expression, mineralization, and β-catenin activation in MC with a much lesser of an effect on NX-MC. YAP/TAZ inhibition also resulted in a reciprocal increase in Smad1/5 phosphorylation as well as BMP2 expression. Our results indicate that increasing MC-GAG stiffness induces osteogenic differentiation via the mechanotransduction mediators YAP/TAZ and the canonical Wnt signaling pathway, whereas the canonical BMPR signaling pathway is activated in a manner independent of mechanical cues.

## Introduction

One of the most exciting developments in the understanding of regenerative technologies is the entry of sophisticated regenerative materials inspired by the extracellular matrix (ECM). The specific composition and material properties of the ECM are known to be essential in instruction of cellular proliferation, differentiation, and tissue organization in both normal physiology as well as pathology (*1*). Thus, ECM-inspired materials have the potential serve as a method for organizing specific regenerative processes. Understanding the exact cell biological consequences of specific properties allows for fine-tuning of such materials and is now one of the frontiers in the development of regenerative technologies.

A primary method for which cells interact with the surrounding extracellular matrix is via integrin interactions at focal adhesions which initiate intracellular changes in the cytoarchitecture via formation of contractile actin-myosin filaments or stress fibers (*2*, *3*). These interactions not only allow for cellular adhesion and spreading, integrin binding also initiates intracellular transduction of signals with the macromolecular assembly and activation of signaling proteins such as focal adhesion kinase (FAK) and the extracellular signal-regulated kinases (ERKs) (*4*, *5*). Both events trigger the activity of the small guanosine triphosphatase (GTPase) Rho (*6*, *7*), which is necessary for the activation of two transcription factors named Yes-associated protein (YAP) and its paralog transcription activator with PDZ-binding motif (TAZ). Both YAP and TAZ are essential to the intracellular transmission of mechanical signals and required for mesenchymal stem cell differentiation induced by stiffness (*8*, *9*).

With respect to the differentiation of progenitor cells such as primary human mesenchymal stem cells (hMSCs), increased stiffness has been found to induce osteogenic differentiation in a YAP/TAZ-dependent manner (*9*). However, the intersection between osteogenic differentiation and mechanical signals is likely complex. One of the best characterized direct relationships is the observation that YAP/TAZ is part of the β-catenin destruction complex in the cytoplasm (*10*). The Wnt (Wingless/integrated) growth factors are important in development and tissue regeneration. Similar to BMPs, Wnt ligands can signal via both canonical (β-catenin-dependent) and non-canonical (β-catenin-independent) pathways (*11*). Negative regulation of the canonical pathway occurs via a cytosolic β-catenin destruction complex assembled on the scaffold protein Axin which interacts with adenomatous polyposis coli (APC), glycogen synthase kinase-3 (GSK3), and casein kinase 1 (CK1) resulting in the phosphorylation of β-catenin by GSK3, ubiquitination by the ubiquitin ligase β-transducin repeats containing protein (β-TrCP), and, ultimately, proteasomal degradation (*11*). With respect to mechanotransduction mediators, YAP and TAZ are binding partners and critical mediators of the β-catenin destruction complex (*10*). In the absence of Wnt activation, YAP and TAZ are required for recruitment of β-TrCP to the destruction complex. In the presence of Wnt activation, YAP, TAZ, and β-catenin are all liberated from the complex, thereby allowing for nuclear translocation and transactivation.

Within the realm of calvarial regeneration, we have previously described the ability of an ECM-inspired nanoparticulate mineralized collagen glycosaminoglycan (MC-GAG) material to induce osteogenic differentiation of primary, bone marrow-derived hMSCs in the absence of exogenous growth factors via autogenously activating the canonical bone morphogenetic protein receptor (BMPR) signaling pathway via phosphorylation of small mothers against decapentaplegic-1/5 (Smad1/5) (*12–18*). More importantly, MC-GAG also induced calvarial regeneration as a progenitor cell-free, exogenous growth factor-free material in rabbits in a manner that approached 60% of the stiffness and strength of the outer cortex of native calvarium (*15*). Our previous work illuminated that the changes in material properties secondary to incorporation of nanoparticulate mineral content was key in the osteogenic properties of MC-GAG as a non-mineralized collagen glycosaminoglycan (Col-GAG) scaffold, fabricated in the same manner, did not demonstrate significant osteogenic abilities (*12*, *13*, *15*, *18*). Given that one of the major differences in MC-GAG and Col-GAG is the stiffness of the material, our current work evaluates the contribution of stiffness and activation of mechanotransduction pathways on the osteogenic abilities of MC-GAG.

## Results

### Stiffness increases the expression of αv, α5, and β1 integrins, phosphorylation of FAK, and YAP/TAZ expression

As the stiffness of the ECM is known to transduce intracellularly via the integrin/FAK/Rho/YAP/TAZ axis, we evaluated the differences between MC-GAG with different rigidities with respect to this axis. Hydrated uncrosslinked (NX-MC) and EDC-crosslinked (MC) MC-GAG have been previously reported (*19*). As NX-MC and MC are fabricated in an identical manner with an identical chemical composition, this study seeks to elucidate the specific contribution of the difference in modulus on hMSC differentiation.

Expression of integrin subunits known to be involved with osteogenic differentiation were evaluated using QPCR at day 3 and 7 of culture (Figure 1A-E). αv (ITGAV), α5 (ITGA5), and β1 (ITGB1) expression were all found to be elevated in MC compared to NX-MC. These differences in gene expression were noted at day 7 of culture. In contrast, α2 (ITGA2) and β3 (ITGB3) were, respectively, downregulated or equivalent in expression patterns between the two scaffolds.

**Figure 1.**
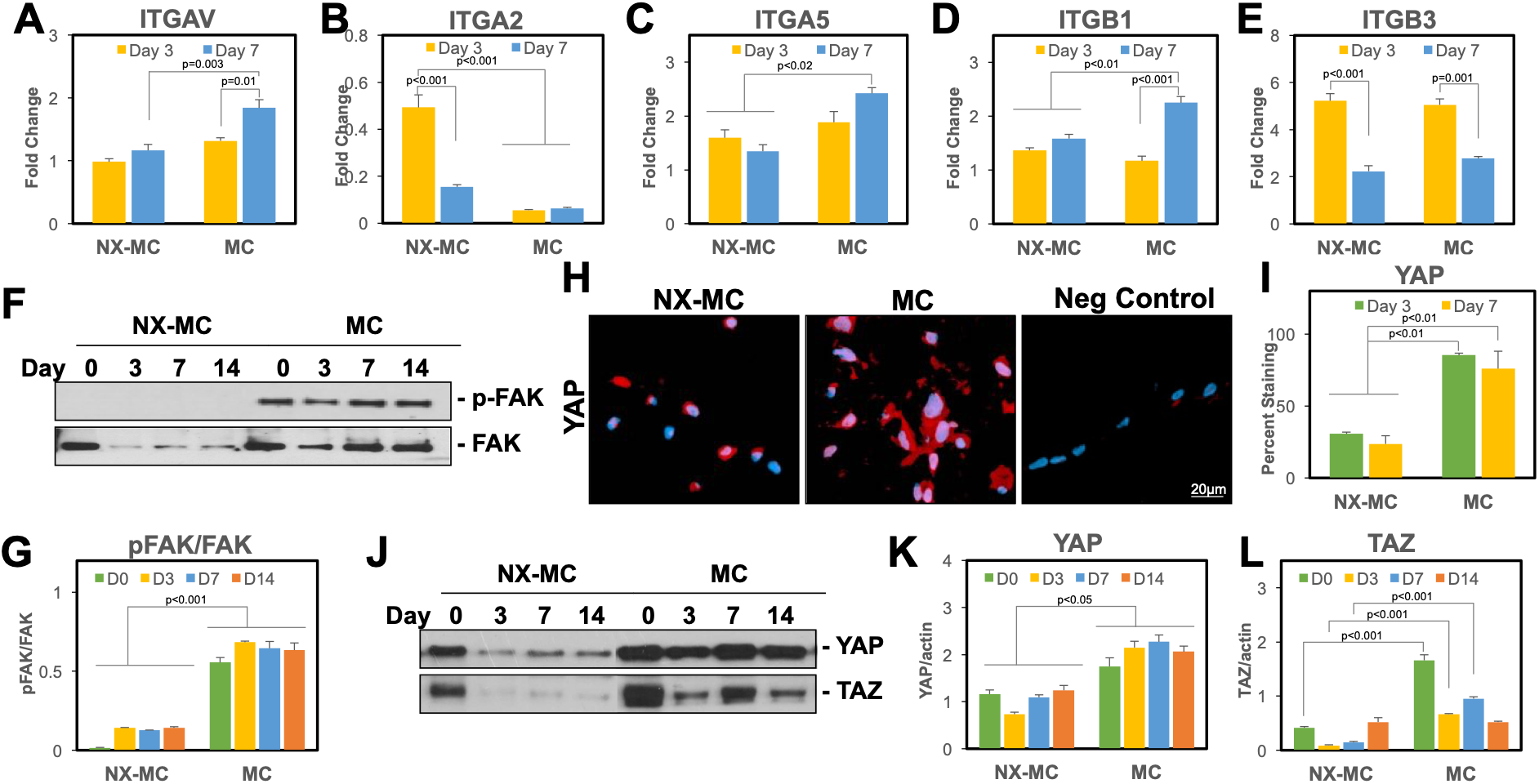
Stiffness increases expression of the αv, α5, and β1 integrin subunits, phosphorylation of FAK, and YAP/TAZ expression. QPCR of primary hMSCs cultured on NX-MC or MC for 3 or 7 days in osteogenic differentiation medium for the (**A**) αv, (**B**) α2, (**C**) α5, (**D**) β1, and (**E**) β3 integrin subunits (n=3-4). Western blot of primary hMSCs cultured on NX-MC or MC for 0, 3, 7, or 14 days in osteogenic differentiation medium for (**F**) phosphorylated FAK (p-FAK) and total FAK or (**J**) YAP and TAZ. Densitometric quantification of western blot analyses demonstrating relative protein amounts of p-FAK to total FAK (**G**), YAP to β-actin (**K**), and TAZ to β-actin (**L**). Merged representative immunofluorescent images of YAP and Dapi costaining (**H**) and quantification (**I**) of YAP staining of primary hMSCs cultured on NX-MC or MC for 3 or 7 days. Bars represent means, errors bars represent SE. Significant posthoc comparisons following ANOVA indicated with p values.

We next evaluated the phosphorylation of FAK, a known key mediator of integrin activated intracellular signals (Figure 1F-G). On western blot analysis, phosphorylation of FAK was significantly elevated on MC compared to NX-MC. Densitometric analyses to quantify the relative ratios of phosphorylated to total FAK revealed that, for comparisons at all timepoints, hMSCs cultured on MC expressed a significantly higher ratio compared to NX-MC (p<0.001 for all comparisons) (Figure 1G).

As key downstream intracellular mediators of mechanotransduction, we next evaluated YAP and TAZ expression (Figure 1H-L). Immunofluorescent staining of YAP was significantly higher in hMSCs cultured on MC versus NX-MC at both day 3 and 7 of culture qualitatively (Figure 1H) as well as quantitatively (Figure 1I). Of note, a significant difference in the morphology of cells was seen in culture on NX-MC versus MC with cells on MC demonstrating more cell spreading whereas cells cultured on NX-MC were small and rounded. On western blot analysis, both YAP and TAZ were increased in hMSCs cultured on MC compared to NX-MC at all timepoints (Figure 1J).

When quantified relative to β-actin expression using densitometry, both YAP and TAZ were significantly increased in MC compared to NX-MC. However, while YAP was elevated at all timepoints on MC, TAZ appeared to be elevated primarily at early timepoints (day 0, 3, and 7) with a reduction at day 14 resulting in equivalent expression to NX-MC.

### Stiffness increases expression of osteogenic genes and Runx2 expression without sustained differences in the activation of the canonical BMPR signaling pathway

To understand the impact of stiffness on the osteogenic capabilities of MC-GAG, we compared the differences in osteogenic gene expression of NX-MC and MC using QPCR (Figure 2A-C). For both NX-MC and MC, expression of the early osteogenic gene ALP increased from day 3 to day 7 in osteogenic differentiation media. While ALP was expressed in higher quantities in MC compared to NX-MC at day 3 (p=0.02), this difference was no longer significant at day 7. For COL1A1 and RUNX2 expression, both genes were increased from day 3 to day 7 only in MC while no significant differences were found in NX-MC over time. For both genes, expression on MC at day 7 was significantly higher compared to NX-MC at either day 3 or day 7 timepoints (p<0.001 for both).

**Figure 2.**
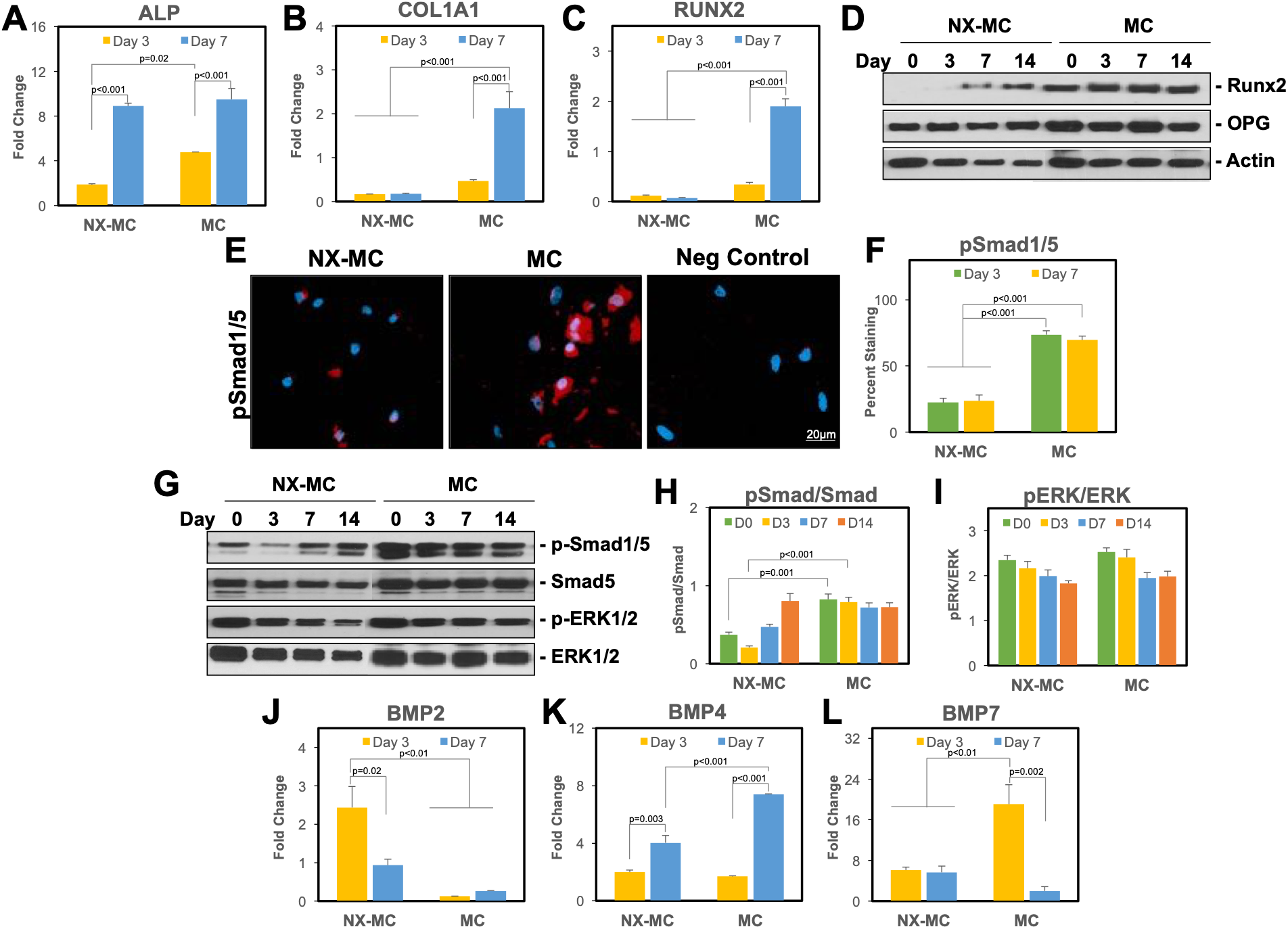
Stiffness increases expression of osteogenic genes and activation of the canonical BMP receptor and Wnt signaling pathways. QPCR of primary hMSCs cultured on NX-MC or MC for 3 or 7 days in osteogenic differentiation medium for (**A**) ALP, (**B**) COL1A1, (**C**) RUNX2, (**J**) BMP2, (**K**) BMP4, and (**L**) BMP7 (n=3). Merged representative immunofluorescent images of phosphorylated Smad1/5 (p-Smad1/5) and Dapi costaining (**E**) and quantification (**F**) of p-Smad1/5 staining of primary hMSCs cultured on NX-MC or MC for 3 or 7 days. Western blot of primary hMSCs cultured on NX-MC or MC for 0, 3, 7, or 14 days in osteogenic differentiation medium for (**D**) Runx2, OPG, and β-actin or (**G**) p-Smad1/5, total Smad5, p-ERK1/2, and total ERK1/2. Densitometric quantification of western blot analyses demonstrating relative protein amounts of p-Smad1/5 to total Smad5 (**H**) and p-ERK1/2 to total ERK1/2 (**I**). Bars represent means, errors bars represent SE. Significant posthoc comparisons following ANOVA indicated with p values.

To evaluate whether the differences in RUNX2 gene expression correlated to protein expression, western blot analysis was performed on primary hMSCs cultured on NX-MC or MC for 0, 3, 7, and 14 days (Figure 2D). Similar to the QPCR results, NX-MC demonstrated less Runx2 protein expression compared to MC at all timepoints. As a comparison, OPG and β-actin protein expression were largely consistent on either material over the course of differentiation. In combination, these data suggested that increasing stiffness with crosslinking MC-GAG increased expression of osteogenic genes and proteins.

Next, we evaluated the major intracellular signaling pathways known to be involved in MC-GAG-mediated as well as stiffness-induced osteogenic differentiation. Previously, our group demonstrated that the canonical BMP receptor signaling pathway was essential for MC-GAG-mediated osteogenic differentiation (*13*, *15*, *18*). Thus, we first evaluated the differences between NX-MC and MC in the phosphorylation of Smad1/5. Immunofluorescent staining of p-Smad1/5 was significantly higher in hMSCs cultured on MC versus NX-MC at both day 3 and 7 of culture qualitatively (Figure 2E) as well as quantitatively (Figure 2F). Representative images demonstrated that the staining of p-Smad1/5 appeared to be both present in the cytoplasm as well as the nucleus.

On western blot analysis, similar to Runx2 expression, phosphorylated Smad1/5 was increased in hMSCs cultured on MC compared to NX-MC at all timepoints (Figure 2G). However, total Smad5 protein also appeared to be expressed in higher quantities in MC compared to NX-MC. Due to the differences in total Smad5 protein, we quantified the western blot bands using densitometric analysis and evaluated the differences in relative amounts of phosphorylated to total protein (Figure 2H). While MC demonstrated a higher p-Smad1/5 to total Smad5 ratio on days 0 and 3 of culture (p<0.001), this difference was no longer significant at later timepoints. Our previous work demonstrated that ERK1/2 phosphorylation (p-ERK1/2) was dispensible to MC-GAG-mediated osteogenic differentiation (*13*, *15*, *18*). Thus, for comparison, we evaluated p-ERK1/2 expression (Figure 2G). Unlike p-Smad1/5, p-ERK1/2 did not demonstrate significant differences between NX-MC and MC, albeit total ERK1/2 was also increased overall in MC. When we quantitatively evaluated the p-ERK1/2 to total ERK1/2 ratio using densitometric analyses, no significant differences were found in the relative expression of p-ERK1/2 to total ERK1/2.

To elucidate the potential reasons for the differences in the activation of Smad1/5 phosphorylation, we evaluated the expression of BMP ligands (Figure 2J-L). While BMP2 expression was downregulated in MC compared to NX-MC, both BMP4 and 7 were upregulated in hMSCs cultured on MC. BMP7 demonstrated an early upregulation at day 3 while BMP4 was upregulated at day 7 on MC compared to NX-MC.

In combination, these data suggest that both NX-MC and MC result in significant activation of Smad1/5, albeit correlating to the upregulation of different BMP ligands. Over time, NX-MC and MC do not significantly differ in the relative expression of p-Smad1/5 to total Smad5, in contrast to the increase in osteogenic gene expression found in MC. Thus, activation of other osteogenic pathways likely distinguishes MC from NX-MC.

### Stiffness activates the canonical Wnt signaling pathway

While the literature is replete with reports on regulatory effects of YAP/TAZ on a variety of cellular processes, one of the best known interactions is the crosstalk of YAP/TAZ with the canonical Wnt/β-catenin signaling pathway (*20*). To evaluate whether the differences in NX-MC and MC correlated to β-catenin activation, we evaluated the expression of the active, non-phosphorylated form of β-catenin (non-p-β-catenin) (Figure 3). Immunofluorescent staining at 3 and 7 days of culture demonstrated that primary hMSCs on MC displayed more non-p-β-catenin compared to NX-MC. When the staining was quantified, the increased non-p-β-catenin in primary hMSCs cultured on MC was found to be statistically significant at both 3 (p<0.01) and 7 days (p<0.001) of culture compared to either timepoint on NX-MC (Figure 3B).

**Figure 3.**
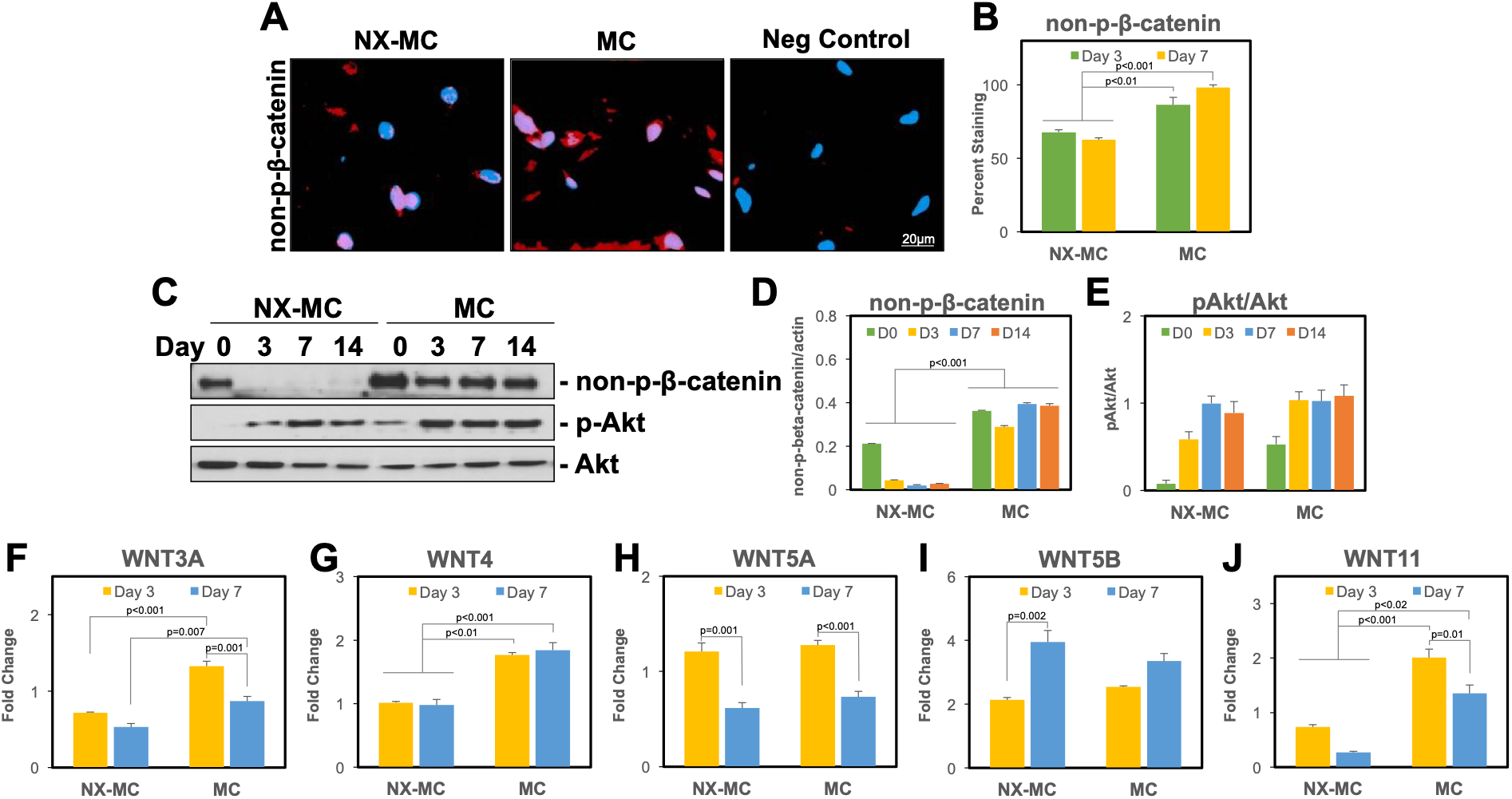
Stiffness activates the canonical Wnt signaling pathway on crosslinked MC-GAG. Merged representative immunofluorescent images of non-phosphorylated-β-catenin (non-p-β-catenin) and Dapi costaining (**A**) and quantification (**B**) of non-p-β-catenin of primary hMSCs cultured on NX-MC or MC for 3 or 7 days. Western blot of primary hMSCs cultured on NX-MC or MC for 0, 3, 7, or 14 days in osteogenic differentiation medium for (**C**) non-p-β-catenin, phosphorylated Akt (p-Akt), and total Akt. Densitometric quantification of western blot analyses demonstrating relative protein amounts of non-p-β-catenin to actin (**D**) and p-Akt to total Akt (**E**). QPCR of primary hMSCs cultured on NX-MC or MC for 3 or 7 days in osteogenic differentiation medium for the (**F**) WNT3A, (**G**) WNT4, (**H**) WNT5A, (**I**) WNT5B, and (**J**) WNT11 (n=3). Bars represent means, errors bars represent SE. Significant posthoc comparisons following ANOVA indicated with p values.

Western blot analyses were also performed to compare the expression of non-p-β-catenin expression over 14 days of culture on NX-MC and MC (Figure 3C). Similar to the expression of p-FAK, YAP, and TAZ, hMSCs cultured on MC demonstrated high expression of non-p-β-catenin. As a comparison, p-Akt and total Akt did not show any significant differences in expression. Densitometric quantification of non-p-β-catenin relative to β-actin expression confirmed that hMSCs cultured on MC were significantly higher than NX-MC at every single timepoint (p<0.001) (Figure 3D). Again, in contrast, activation of Akt relative to total Akt was not statistically significantly different at any timepoint compared between MC versus NX-MC.

One of the hallmarks of YAP/TAZ activation is the amplification of signals via a positive feedback loop (*21*). In the setting of β-catenin activation, YAP and TAZ clearly have direct effects on liberating β-catenin from the destruction complex but also may act indirectly to further augment β-catenin activation by increasing Wnt ligand expression. Thus, we next evaluated whether increases in Wnt ligand transcription could be detected (Figure 3F-J). hMSCs cultured on NX-MC and MC and differentiated for 3 or 7 days were subjected to QPCR for a panel of Wnt ligands. Among the ligands, WNT5A and WNT5B demonstrated no significant differences between NX-MC and MC. WNT3A, WNT4, and WNT11 displayed a significant upregulation in MC compared to NX-MC at both day 3 and 7 of culture in a statistically significant fashion. In combination, these data suggest that MC scaffolds upregulate the canonical Wnt signaling pathway in a manner that correlated to an upregulation of WNT3A, WNT4 and WNT11 expression.

### Inhibition of Rho GTPase or F-actin polymerization abrogates differences in YAP/TAZ expression, FAK phosphorylation, and αv and α5 integrin expression on MC versus NX-MC

To understand the necessity of YAP/TAZ mechanotransduction in the osteogenic capabilities of MC, we next inhibited the small GTPase Rho and actin polymerization using C3 transferase and latrunculin A (LA), respectively, as both are known to be required for YAP/TAZ activation (*9*). Primary hMSCs differentiated on NX-MC and MC were untreated or treated with LA or C3 and western blotted for YAP and TAZ (Figure 4A-C). Again, expression of both YAP and TAZ were found to be significantly higher in hMSCs differentiated on MC compared to NX-MC both qualitatively (Figure 4A) and quantitatively (p<0.001) (Figure 4B-C). For both NX-MC and MC, LA demonstrated a stronger inhibition of YAP and TAZ compared to C3. Quantitatively, both inhibitors significantly reduced the expression of both YAP and TAZ for MC scaffolds. On NX-MC, LA significantly reduced YAP (p<0.001) and TAZ (p=0.02) expression, while the reduction by C3 did not reach statistical significance compared to untreated control. To confirm that inhibition of YAP and TAZ expression correlated to reduction in expression of known downstream targets, we evaluated the expression of the best characterized YAP/TAZ target, CTGF, using QPCR (Figure 4D). For hMSCs cultured on both NX-MC and MC, CTGF expression was reduced in the presence of either LA or C3.

**Figure 4.**
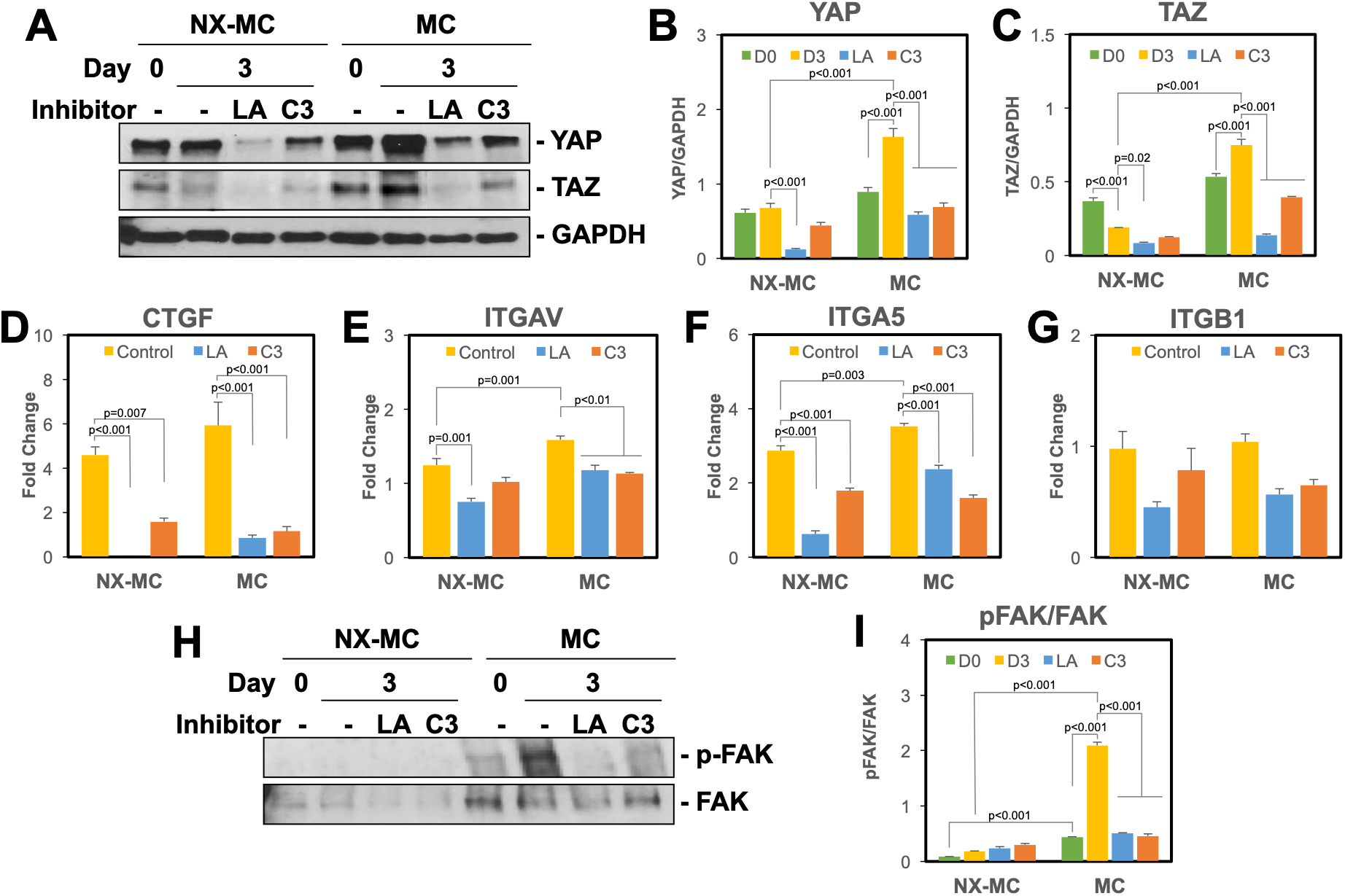
Inhibition of YAP and TAZ activation via latrunculin A and C3 also inhibits expression of α5 integrin subunit and phosphorylation of FAK on crosslinked MC-GAG. Western blot of primary hMSCs cultured on NX-MC or MC for 0 or 3 days in osteogenic differentiation medium untreated, treated with 0.5 μM of latrunculin A (LA), or 3 μg/mL of C3 for (**A**) YAP, TAZ, and GAPDH or (**H**) p-FAK and total FAK. Densitometric quantification of western blot analyses demonstrating relative protein amounts of YAP to GAPDH (**B**), TAZ to GAPDH (**C**), and p-FAK to FAK (**I**). QPCR of primary hMSCs cultured on NX-MC or MC for 3 days in osteogenic differentiation medium untreated, treated with 0.5 μM of LA, or treated with 3 μg/mL of C3 for the (**D**) CTGF, (**E**) ITGAV, (**F**) ITGA5, and (**G**) ITGB1 (n=3-4). Bars represent means, errors bars represent SE. Significant posthoc comparisons following ANOVA indicated with p values.

While mechanotransduction via YAP and TAZ is frequently considered to be downstream of focal adhesions, YAP and TAZ knockdown are also known to disrupt the integrity of focal adhesions partially by downregulating expression of multiple integrin subunits suggesting bidirectional regulation (*22*). As we noted that ITGAV, ITGA5, and ITGB1 are upregulated in hMSCs differentiated on MC (Figure 1A-E), we evaluated the expression of these subunits in the presence of LA and C3 treatment (Figure 4E-G). Both ITGAV and ITGA5 were downregulated by LA and C3 treatment for MC scaffolds (p<0.01 and p<0.001, respectively with either inhibitor). For ITGAV, downregulation on NX-MC was also found in the presence of LA (p<0.001) but not C3 (Figure 1E), consistent with the weaker inhibition of YAP and TAZ found with C3 treatment. While ITGA5 was downregulated in the presence of either inhibitors on NX-MC (p<0.001), the reduction by C3 treatment was less relative to LA. In contrast ITGB1 was unaffected by either inhibitor. These data suggest that both ITGAV and ITGA5 expression are regulated by YAP and TAZ on MC in a manner that is dependent upon both Rho GTPase and actin polymerization. On NX-MC, while YAP and TAZ are likely to still play a role in mechanotransduction, Rho inhibition appears to have less of an effect on YAP and TAZ expression as well as expression of ITGAV and ITGA5.

To understand the downstream effects of changes in integrin expression on signal transduction, we evaluated the expression of phosphorylated FAK in western blot analysis in the presence of LA and C3 (Figure 4H-I). Again, the ratio of p-FAK to total FAK was quantitatively less in NX-MC scaffolds at day 0 as well as day 3 (p<0.001). In NX-MC scaffolds, the addition of LA and C3 did not significantly affect the amount of phosphorylated to total FAK expression. In contrast, both LA and C3 treatment significantly reduced the amount of p-FAK to FAK on MC scaffolds compared to untreated controls (p<0.001).

### Osteogenic gene expression and mineralization is reduced by inhibition of Rho GTPase or F-actin polymerization on MC

To understand the necessity of YAP and TAZ on the differences in osteogenic gene expression between MC compared to NX-MC, we evaluated the expression of ALP, COL1A1, and RUNX2 untreated and treated with LA or C3 (Figure 5A-C). While MC expressed significantly more ALP compared to NX-MC in untreated cells in QPCR, LA or C3 treatment significantly reduced ALP expression on MC (p=0.004 and p=0.006, respectively) to a level comparable to untreated NX-MC (Figure 5A). ALP expression on NX-MC was also reduced by LA but did not demonstrate any statistically significant differences in the presence of C3. Similarly, COL1A1 expression on MC was reduced in the presence of LA and C3 compared to untreated MC (p<0.001 for both inhibitors) to levels comparable to that of NX-MC. No significant differences were notable on NX-MC for COL1A1 expression in the presence of the inhibitors (Figure 5B). Lastly, similar findings were noted for RUNX2 gene expression on MC with significant reduction in the presence of LA or C3 (p=0.002 and p=0.003, respectively) compared to the untreated MC. The reduced levels were comparable to NX-MC. Of note, no differences in RUNX2 gene expression could be elicited by LA or C3 treatment on NX-MC scaffolds (Figure 5C). Interestingly, despite the reduction in RUNX2 expression in the presence of the inhibitors on MC scaffolds, RUNX2 expression was still upregulated approximately two-fold.

**Figure 5.**
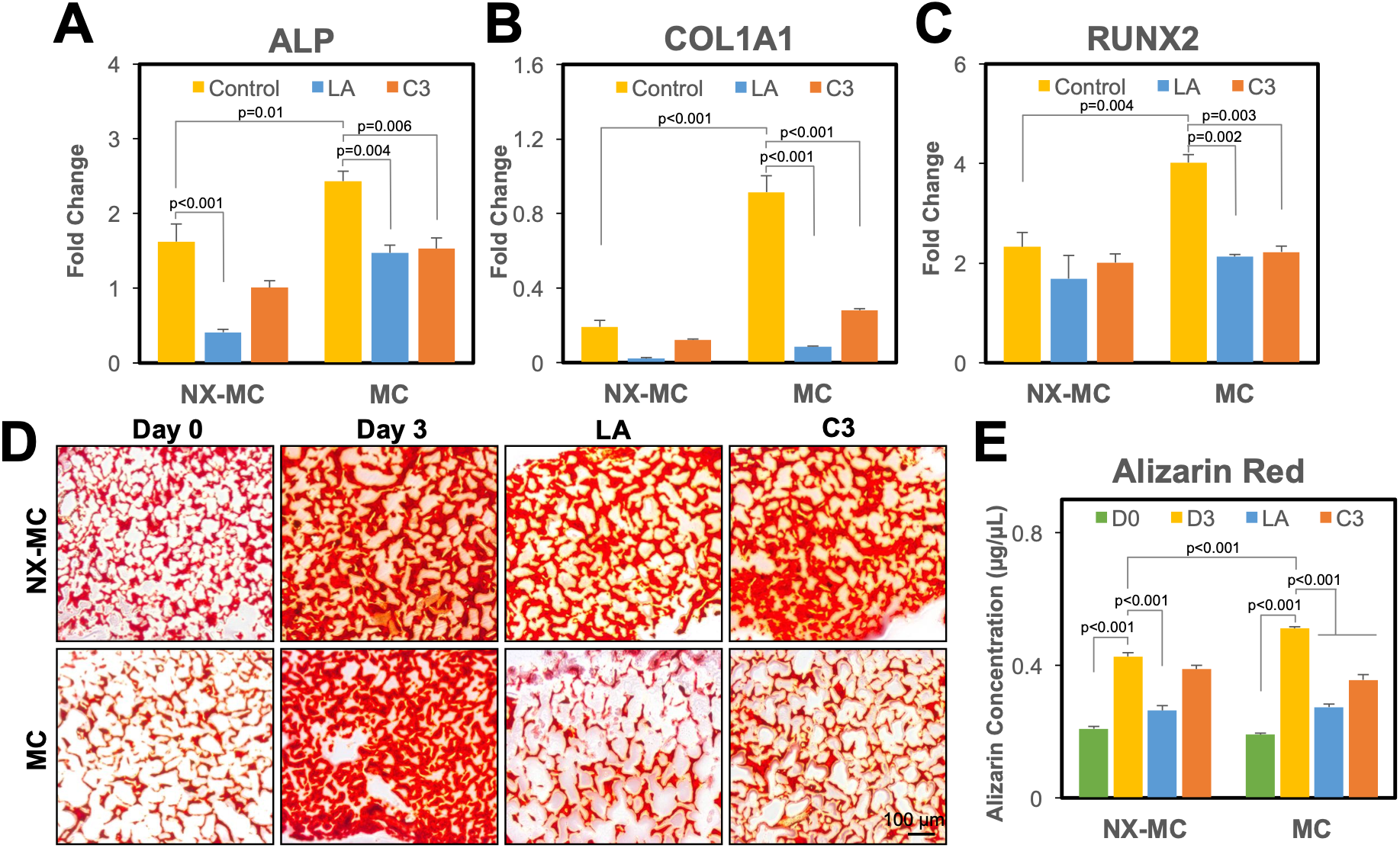
Latrunculin A and C3 inhibits osteogenic gene expression on MC but not NX-MC. QPCR of primary hMSCs cultured on NX-MC or MC for 0 or 3 days in osteogenic differentiation medium untreated, treated with 0.5 μM of LA, or treated with 3 μg/mL of C3 for (**A**) ALP, (**B**) COL1A1, and (**C**) RUNX2. (**D**) Representative images and (**E**) quantitative Alizarin red staining of 5 μm sections of hMSCs cultured on NX-MC or MC for 0 or 3 days in osteogenic differentiation medium untreated, treated with 0.5 μM of LA, or treated with 3 μg/mL of C3 (n=4-5). Bars represent means, errors bars represent SE. Significant posthoc comparisons following ANOVA indicated with p values.

The requirement of YAP and TAZ activation was also evaluated with respect to mineralization of hMSCs using Alizarin red staining (Figure 5D-E). Compared to the negative control (day 0), an increase in mineralization was detected after 3 days of culture for both NX-MC and MC (p<0.001). Consistent with the osteogenic gene expression, MC demonstrated an increased amount of mineralization compared to NX-MC (p<0.001). When treated with LA, both NX-MC and MC were reduced in mineralization compared to the respective untreated scaffolds (p<0.001). C3, again, showed a weaker but significant inhibition of Alizarin red staining on MC (p<0.001) whereas no differences were seen on NX-MC. In combination, these data demonstrate that the increased osteogenic gene expression and mineralization in MC compared to NX-MC could be abrogated with the inhibition of YAP and TAZ activation via LA or C3 treatment. Osteogenic gene expression and mineralization on NX-MC are significantly less affected by either LA or C3.

### Inhibition of Rho GTPase or F-actin polymerization reduces active β-catenin with a compensatory increase in p-Smad1/5 and BMP2 expression

As inhibition of YAP/TAZ via latrunculin A or C3 transferase abrogates the differences in the osteogenic gene expression of MC compared to NX-MC, we next evaluated the activation of the osteogenic signaling pathways. Similar to QPCR, the elevation of Runx2 protein expression on MC was reduced in the presence of LA whereas NX-MC demonstrated no significant differences (Figure 6A-B). While C3 also demonstrated a reduction in Runx2 protein, the reduction did not reach statistical significance on densitometric quantification. For comparison, no significant differences were noted in OPG protein expression.

**Figure 6.**
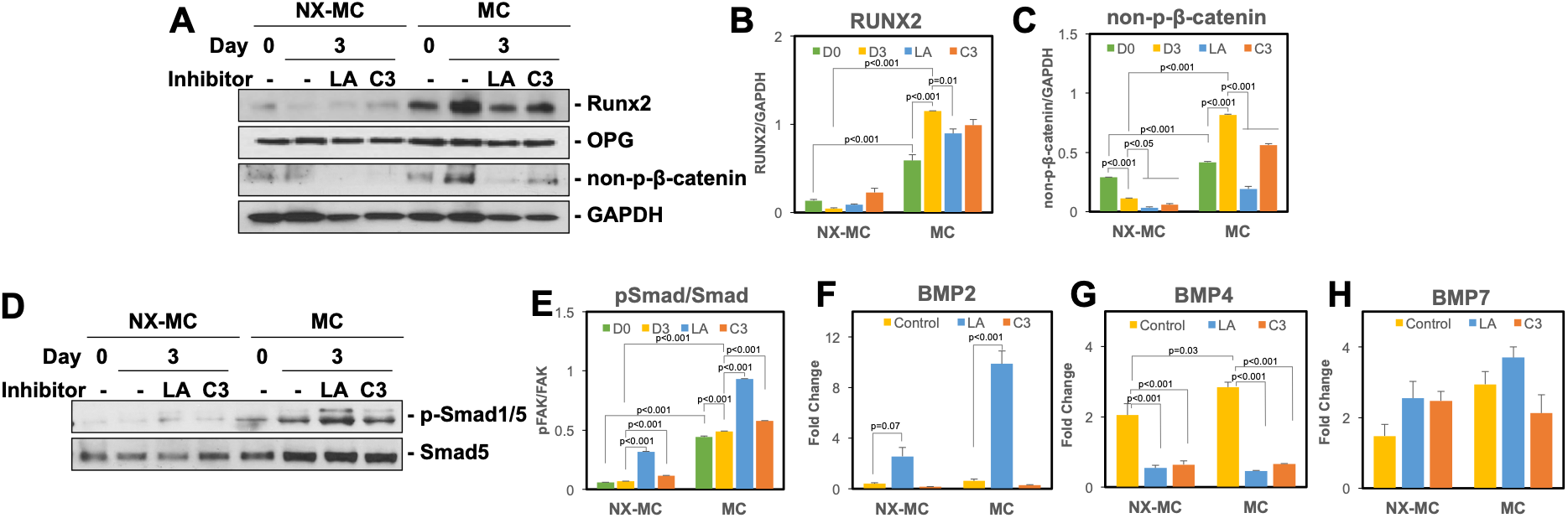
Inhibition of Rho GTPase or F-actin polymerization reduces active β-catenin with a compensatory increase in p-Smad1/5 and BMP2 expression. Western blot of primary hMSCs cultured on NX-MC or MC for 0 or 3 days in osteogenic differentiation medium untreated, treated with 0.5 μM of LA, or treated with 3 μg/mL of C3 for (**A**) Runx2, OPG, non-p-β-catenin, and GAPDH or (**D**) p-Smad1/5 and Smad5. Densitometric quantification of western blot analyses demonstrating relative protein amounts of (**B**) Runx2 to GAPDH, (**C**) non-p-β-catenin to GAPDH, or (**E**) p-Smad1/5 to Smad5. QPCR of primary hMSCs cultured on NX-MC or MC for 0 or 3 days in osteogenic differentiation medium untreated, treated with 0.5 μM of LA, or treated with 3 μg/mL of C3 for (**F**) BMP2, (**G**) BMP4, and (**H**) BMP7. Bars represent means, errors bars represent SE. Significant posthoc comparisons following ANOVA indicated with p values.

To understand the necessity of YAP/TAZ activation for the differences in the activation of the canonical Wnt pathway in MC versus NX-MC, we next evaluated the expression of non-p-β-catenin in the absence and presence of LA and C3 (Figure 6A, C). As demonstrated previously (Figure 3C), non-p-β-catenin was highly activated in MC compared to NX-MC (p<0.001). In the presence of either LA or C3, both NX-MC and MC demonstrated a significant decrease in non-p-β-catenin expression (p<0.05 and p<0.001, respectively). Again, the magnitude of the decrease in expression on MC scaffolds was larger in the presence of LA compared to C3, corresponding to the strength of inhibition of LA compared to C3 of YAP, TAZ, and the osteogenic genes (Figures 4 and 5).

Interestingly, while both LA and C3 demonstrated dramatic effects on YAP, TAZ, and non-p-β-catenin expression, the effects on Runx2 gene and protein expression were partial, suggesting potentially that compensatory activation of other osteogenic pathways may be occurring. Thus, we evaluated activation of p-Smad1/5 in the canonical BMPR signaling pathway (Figure 6D-E). Unlike non-p-β-catenin, treatment with LA or C3 resulted in an increase on p-Smad1/5 compared to untreated NX-MC or MC scaffolds (p<0.001 for all comparisons). When transcription of the BMP ligands was evaluated, BMP2 also demonstrated a reciprocal increase in transcription in the presence of LA in a significant manner on MC (Figure 6F). While expression of BMP2 was significantly lower on NX-MC scaffolds, a trend towards increased BMP2 expression was also found in the presence of LA (p=0.07). In contrast, BMP4 expression patterns paralleled that of non-p-β-catenin with decreases in the presence of LA or C3 for either scaffold and BMP7 exhibited no differences in expression in the presence of either inhibitor on either scaffold.

## Discussion

In this work, we investigated the contribution of mechanotransduction in the osteogenic capabilities of MC-GAG. By comparing the same material in a non-crosslinked versus crosslinked form, we demonstrated that the increase in stiffness from crosslinking correlated to an increase in the expression of a specific integrins (αv, α5, and β1), upregulation of FAK phosphorylation, and an increase in the expression of the major mechanotransduction transcription factors YAP and TAZ in primary, bone marrow-derived human mesenchymal stem cells. Crosslinked scaffolds demonstrated an increase in expression of early osteogenic genes (ALP and COL1A1) as well as an increase in both gene and protein expression of RUNX2. Given that we previously described the autogenous activation of the canonical BMPR signaling pathway by crosslinked MC-GAG, we evaluated the expression of phosphorylated Smad1/5 in relationship to total Smad5 between the two materials. While the p-Smad1/5 to Smad5 ratio was elevated in crosslinked scaffolds at the earliest timepoints (day 0 and 3), later timepoints did not show any statistically significant differences between the two materials suggesting that both materials were equivalent in the activation of the canonical BMPR signaling pathway. Interestingly, there was a difference in the type of BMP ligands expressed with the non-crosslinked version expressing more BMP2 early whereas the crosslinked scaffold expressed more BMP7 early followed by an increase in BMP4 expression. These data suggested that other osteogenic pathways were likely responsible for the differences between non-crosslinked and crosslinked materials. As one of the best characterized osteogenic pathways downstream of mechanical signals, we evaluated the canonical Wnt pathway between the two materials. Similar to the expression of phosphorylated FAK, YAP, and TAZ, the expression of active, non-phosphorylated β-catenin was highly upregulated in crosslinked scaffolds. This expression was correlated to an increase in expression of WNT3A, WNT4, and WNT11. To determine the necessity of the mechanotransduction pathways, we then employed two inhibitors known to prevent the YAP and TAZ activation via inhibition of Rho GTPase (C3 transferase) and inhibition of actin polymerization (latrunculin A). For both inhibitors, reductions in expression of YAP, TAZ, αv integrin, and α5 integrin were demonstrated for either scaffold. Interestingly, for the expression of osteogenic genes ALP, COL1A1, and RUNX2, C3 and latrunculin A primarily reduced expression in the crosslinked scaffolds, while hMSCs cultured on the non-crosslinked materials were largely unaffected. The reduction in expression of the osteogenic genes was paralleled by a reduction in the expression of non-phosphorylated β-catenin in the presence of the inhibitors on the crosslinked scaffolds. However, the reduction in active β-catenin coincided with a reciprocal increase in Smad1/5 phosphorylation and BMP2 expression. These data suggest several conclusions: 1. Holding chemical composition and fabrication equal, mechanical signals from MC-GAG trigger osteogenic differentiation of hMSCs; 2. Mechanotransduction-mediated osteogenic signals activate β-catenin in the canonical Wnt signaling pathway on MC-GAG without affecting Smad1/5 phosphorylation; 3. Inhibition of mechanotransduction reduces osteogenic differentiation but does not eliminate differentiation; 4. Inhibition of mechanotransduction on MC-GAG results in a reciprocal activation of Smad1/5 phosphorylation.

The understanding of cell biology induced by regenerative materials is essential to fine-tuning of materials to maximize intentional cellular consequences such as differentiation while minimizing unintentional consequences such as hyperproliferation or inflammation. Our previous work had demonstrated the MC-GAG induced osteogenic differentiation of primary hMSCs and *in vivo* rabbit calvarial regeneration in a manner that was superior to a non-mineralized version of the material (Col-GAG) and in a manner that did not require growth factor induction. As these findings positioned MC-GAG as a potential materials-only solution for calvarial regeneration, a thorough understanding of the material properties responsible for these observations would benefit material refinement. In comparison to Col-GAG, the addition of nanoparticulate mineral in MC-GAG contributes two major material differences: an increase in stiffness and a depot of calcium and phosphate ions. Our current work isolated the mechanical signal by evaluating MC-GAG at two different stiffnesses and found that the crosslinked material allowed for improved osteogenic differentiation via the canonical Wnt pathway. While these results are useful as a starting point, potentially a range of stiffnesses with varying techniques of crosslinking may be necessary to further refine the material to optimize β-catenin activation. Another interesting observation from these results is that the activation of Smad1/5 phosphorylation was largely independent from the differences in stiffness. As Col-GAG minimally induces p-Smad1/5 expression and the other major difference between MC-GAG and Col-GAG is the presence of calcium and phosphate ions, it is likely that the inorganic ion-mediated signaling pathways may be the trigger for activation of the canonical BMPR signaling pathways (Figure 7). These investigations are under way in our group.

**Figure 7.**
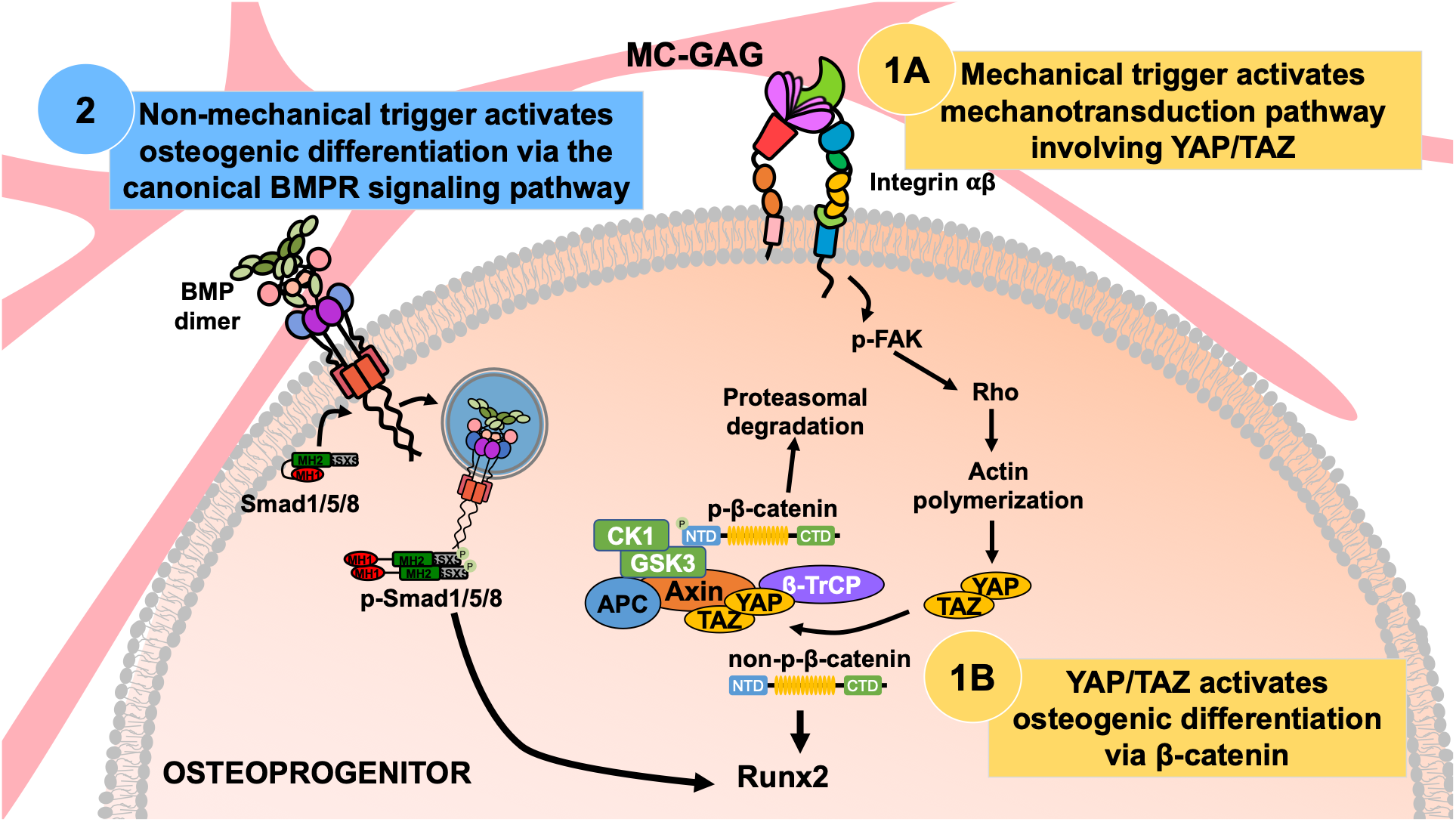
Model of MC-GAG mediated osteogenic differentiation pathways. Mechanical signals, such as stiffness, from MC-GAG contribute to inducing differentiation of osteoprogenitors in a manner that requires activation of YAP/TAZ (1A) resulting in the activation of β-catenin (1B). In contrast, osteogenic differentiation downstream of activation of the canonical BMPR signaling pathway occurs in a YAP/TAZ-independent manner on MC-GAG (2), suggesting that this pathway is activated via a non-mechanical trigger.

While other investigators have also reported the relationship between stiffness of materials and induction of ostegeogenic differentiation, a number of differences are present between other studies and this work. Du and colleagues noted that chondrocytes seeded on polyacrylamide of varying stiffness also demonstrated activation of β-catenin and reduction in cartilage markers in a manner that required integrin and FAK phosphorylation (*20*). Using polyacrylamide coated with fibronectin varying in stiffness from 13-68 kPa, Sun et al also demonstrated an improvement in osteogenic differentiation on stiffer substrates in a manner that required α5β1 integrin signaling resulting in ERK phosphorylation, Akt phosphorylation, and β-catenin activation (*23*). Both the former studies contrast with our work in that activation of Akt and ERK were uninvolved in our results, potentially secondary to cell type differences or differences in the composition of the materials used. The importance of scaffold composition with respect to the identity of ECM proteins was recently evaluated by Stanton and colleagues (*24*). Their major findings were that while nuclear translocation of YAP and osteogenic differentiation was increased in stiffer hydrogels, the type and density of the immobilized ECM protein were relevant such that YAP translocation was primarily associated with collagen I or IV but not laminin or fibronectin. Olivares-Navarrete and colleagues have previously reported that titanium surfaces increased non-canonical Wnt5a signaling and osteogenic differentiation whereas the canonical Wnt3a/β-catenin pathways were not involved (*25*). Within the same group, a followup study showed a contribution for Wnt11, a non-canonical Wnt thought to be a stabilizer of β-catenin, thereby connecting the contributions of both canonical and non-canonical pathways for hMSC differentiation (*26*). The combination of these studies suggest that stiffness is likely only one component of material properties that direct differentiation and that the composition of proteins as well as inorganic substances are important for the variabilities in signaling pathways activated. Thus, individual materials induce significant differences in cell biology that must be characterized if refinement is to be realized.

While not the subject of this study, it is important to comment on the surgical practicality of the mineralized collagen scaffolds studied here. This concern is driven by differences in macroscale vs. microscale mechanical performance criteria that are often in opposition. The macroscale mechanical perfomance of these materials such as its modulus (*E*\*) is primarily influenced by the high porosity (>85%) of the scaffold. The modulus of low-density, open-cell foams, the class of material to which these scaffolds belong, scales as the modulus of the solid material from which the foam is fabricated (*E_s_*) and the relative density (*ρ*\*/*ρ_s_*) of the foam squared. So while individual cells experience a microscale mechanical defined by *E_s_*, the bulk performance of the scaffold *E*\* is dictated by the high porosity necessary to support penetration, metabolic health, and tissue biosynthesis. To address the need to improve macroscale mechanical performance without sacrificing porosity, we have recently reported approaches to integrate a millimeter-scale reinforcing poly (lactic acid) (PLA) frame fabricated via 3D-printing into the mineralized collagen scaffold to form a multi-scale mineralized collagen-PLA composite (*27*). This composite shows significant increased macroscale mechanical performance (Young’s Modulus, ultimate stress, ultimate strain) that is dictated by the PLA frame design, but osteogenic activity dictated by the scaffold. We also demonstrated adaptations to the PLA frame to render the composite able to conformally-fit complex defect margins. So while this study primarily focuses on insights regarding crosslinking-induced changes in microscale mechanical performance (*28*), we have methods to render these scaffolds mechanically competent for surgical use.

The classical paradigm of regenerative technologies consisting of progenitor cell and growth factor delivery on a material has not been realized in the realm of surgical and clinical practicality despite 30 years of research. In skeletal reconstruction, the idea of harvesting stem cells for *ex vivo* expansion and growth is highly impractical in practice as well as in expense as autologous methods for bone grafting or transfer of free-vascularized bone, despite its morbidity, can be achieved. While the idea of growth factor delivery for augmentation of skeletal regeneration has been of great interest, the two FDA-approved growth factors for bone have shown that physiologic dysregulation with supraphysiologic dosages of single factors is not likely to be the ideal strategy for regeneration. One example is the fact that a decreasing usage of BMP-2 in anterior spinal fusion has occurred due to the reported increases in complications compared to traditional, growth factor-free, autologous bone grafting (*29*). In the past decade, we have seen a rise in interdisciplinary collaborations between surgeons, cell biologists, and bioengineers in an effort towards understanding regeneration. One of the themes that has appeared over and over again is that there is not one method or one factor that will ultimately recapitulate biology in a manner that is adequate, safe, or practical. With these ideas in mind, creation of microenvironments that allow for influx of endogenous progenitor cells that provide a framework for organization may likely be the closest artificial method for regeneration of single tissue types as well as tissues that are non-essential.

## Materials and Methods

### Fabrication of mineralized collagen scaffolds

MC-GAG scaffolds were prepared using the lyophilization process described previously (*30–32*). Briefly, a suspension of collagen and GAGs were produced by combining microfibrillar, type I collagen (Collagen Matrix, Oakland, NJ) and chondroitin-6-sulfate (Sigma-Aldrich, St. Louis, MO) with calcium salts (calcium nitrate hydrate: Ca(NO_3_)_2_·4H_2_O; calcium hydroxide: Ca(OH)_2_, Sigma-Aldrich, St. Louis, MO) in a solution of phosphoric acid. The suspension was frozen using a constant cooling rate technique (1 °C/min) from room temperature to a final freezing temperature of −10 °C using a freeze dryer (Genesis, VirTis). Following sublimation of the ice phase, scaffolds were sterilized via ethylene oxide and cut into 8 mm disks for culture. NX-MC scaffolds were rehydrated with phosphate buffered saline (PBS) overnight and used for cell culture.

### Chemical crosslinking of mineralized collagen scaffolds

Crosslinking of scaffolds were prepared by rehydrating scaffolds in PBS overnight with 1-ethyl-3-(3- dimethylaminopropyl) carbodiimide (EDAC, Sigma-Aldrich) and N-hydroxysuccinimide (NHS, Sigma Aldrich) at a molar ratio of 5:2:1 EDC:NHS:COOH where COOH represents the amount of collagen in the scaffold as we previously described (*33*). Scaffolds were washed with PBS to remove any of the residual chemical.

### Cell culture

Primary hMSCs (Lonza, Inc., Allendale, NJ) were expanded in Dulbecco’s Modified Eagle’s medium (DMEM, Corning Cellgro, Manassas, VT) supplemented with 10% fetal bovine serum (Atlanta Biologicals, Atlanta, GA), 2 mM L-glutamine (Life Technologies, Carlsbad, CA), 100 IU/mL penicillin/100 μg/mL streptomycin (Life Technologies). After expansion, 3 × 10^5^ hMSCs were seeded onto 8 mm NX-MC and MC scaffolds in growth media. 24 h after seeding, media was switched to osteogenic differentiation media consisting of 10 mM β-glycerophosphate, 50 μg/mL ascorbic acid, and 0.1 μM dexamethasone. NX-MC and MC scaffolds were treated or untreated with 0.5 μM latrunculin A (LA, Santa Cruz Biotechnology, Santa Cruz, CA) or 3 μg/mL C3 transferase (Cytoskeleton, Inc, Denver, CO).

### Quantitative real-time reverse-transcriptase polymerase chain reaction

Total RNA was extracted using the RNeasy kit (Qiagen, Valencia, CA) at 0, 3, and 7 days of culture. Gene sequences were obtained from the National Center for Biotechnology Information gene database and primers were designed (Eurofins Genomics, Louisville, KY, Table 1). Quantitative real time RT-PCR (QPCR) were performed on the Opticon Continuous Fluorescence System (Bio-Rad Laboratories, Inc., Hercules, CA) using the QuantiTect SYBR Green RT-PCR kit (Qiagen). Cycle conditions were as follows: reverse transcription at 50°C (30 min); activation of HotStarTaq DNA polymerase/inactivation of reverse transcriptase at 95°C (15 min); and 45 cycles of 94°C for 15s, 58°C for 30 s, and 72°C for 45 s. Results were analyzed and presented as representative graphs of triplicate experiments.

**Table 1.**
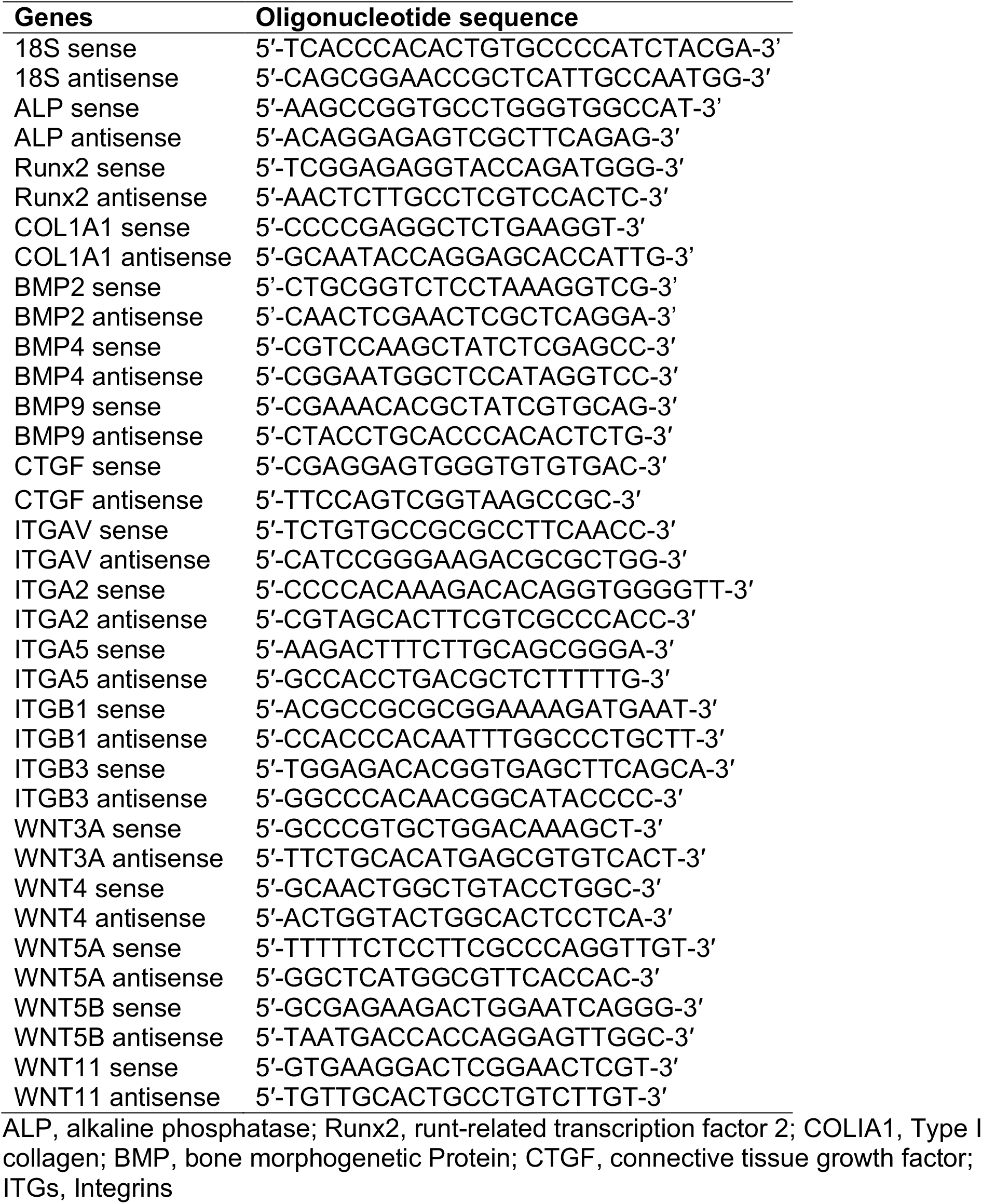
Primer Sequences

### Western blotting

Lysates for western blot analysis were prepared from scaffolds at 0, 3, 7 and 14 days of culture using Phosphosafe Lysis Buffer (Novagen, Madison, WI). Equal amounts of protein lysates (10-20 μg total protein) were subjected to 4–20% SDS-PAGE (Bio-Rad, Hercules, CA). Western analysis was carried out with primary antibodies against phosphorylated Smad1/5 (p-Smad1/5), total Smad5, phosphorylated Akt (p-Akt), total Akt, phosphorylated FAK (p-FAK), total FAK, phosphorylated ERK1/2 (p-ERK1/2), total ERK1/2, OPG, RUNX2, YAP, TAZ, non-phosphorylated-β-catenin (non-p-β-catenin), GAPDH, and β-actin and the respective species-specific horseradish peroxidase-conjugated secondary antibodies (Santa Cruz Biotechnology, Santa Cruz, CA). Detection was performed using an enhanced chemiluminescent substrate (Thermo Scientific, Rockford, IL). All primary antibodies against phosphorylated proteins were obtained from Cell Signaling Technologies (Beverly, MA), and all primary full-length antibodies were obtained from Santa Cruz Biotechnology (Santa Cruz, CA). Imaging was carried out and quantified using Image J (NIH, Bethesda, MD).

### Qualitative and Quantitative Alizarin Red Staining

Histology was performed on scaffolds at 0, 3, 7 days of culture. Scaffolds were fixed in 10% formalin, paraffin-embedded, and sectioned at 4 μm. Following deparaffinization, the sections were stained with Alizarin Red at 10 mg/ml (Cell Signaling Technologies, Beverly, MA). Slides were analyzed qualitatively using a standard microscope and digitally photographed. For quantification, add 10% glacial acetic acid (Fisher Scientific International, Inc., Pittsburgh, MA) to the staining sections for 30 minutes. Absorbance of the incubation medium was measured at 450 mm (Epoch spectrophotometer, BioTek, Winooski, VT).

### Immunofluorescence Microscopy

Scaffolds were fixed in 10% formalin, paraffin-embedded, and sectioned at 4 μm. Following deparaffinization, the sections were treated with 0.5% Triton X-100 (MP Biomedicals, LLC, Santa Ana, CA), blocked with 10% normal goat serum (Jackson ImmunoResearch Laboratories, Inc, West Grove, PA) for 1 hour, and subjected to heat-induced antigen retrieval using sodium citrate buffer(10 mM, pH 6, Fisher Scientific International, Inc., Pittsburgh, MA) at >80°C, for 20 minutes. Sections were then separately incubated with anti-p-Smad1/5 (1:500, Cell Signaling Technologies), anti-Yap (1:200, Santa Cruz Biotechnology, Santa Cruz, CA), anti-non-p-β-catenin (1:1000 Cell Signaling Technologies, Beverly, MA) overnight in 4 °C. After washing, sections were incubated in anti-rabbit or anti-mouse IgG Alexa Fluor Plus 594 (Thermo Fisher, Eugene, OR). Coverslips were mounted with Prolong Gold Antifade Reagent with Dapi (Cell Signaling Technologies, Danvers, MA). Images were captured with Zeiss Axio Observer 3 inverted microscope with ZEN 2.3 Pro software (Zeiss, Oberkochen, Germany).

### Statistical analysis

All statistical analyses were performed using SPSS Version 25 (Chicago, IL) using analyses of variance (ANOVA) followed by post hoc tests under the Tukey criterion. A value of p<0.05 was considered significant.

